# Transcriptome profiling of pathogen-specific CD4 T cells identifies T-cell-intrinsic caspase-1 as an important regulator of Th17 differentiation

**DOI:** 10.1101/452763

**Authors:** Yajing Gao, Krystin Deason, Aakanksha Jain, Ricardo A Irizarry-Caro, Igor Dozmorov, Isabella Rauch, Edward K Wakeland, Chandrashekhar Pasare

## Abstract

Our study revealed that DCs shape distinct pathogen-specific CD4 T cell transcriptome and from which, we discovered an unexpected role for T-cell-intrinsic caspase-1 in promoting Th17 differentiation.

**ABSTRACT:** Dendritic cells (DCs) are critical for priming and differentiation of pathogen-specific CD4 T cells. However, to what extent innate cues from DCs dictate transcriptional changes in T cells leading to effector heterogeneity remains elusive. Here we have used an *in vitro* approach to prime naïve CD4 T cells by DCs stimulated with distinct pathogens. We have found that such pathogen-primed CD4 T cells express unique transcriptional profiles dictated by the nature of the priming pathogen. In contrast to cytokine-polarized Th17 cells that display signatures of terminal differentiation, pathogen-primed Th17 cells maintain a high degree of heterogeneity and plasticity. Further analysis identified caspase-1 as one of the genes upregulated only in pathogen-primed Th17 cells but not in cytokine-polarized Th17 cells. T-cell-intrinsic caspase-1, independent of its function in inflammasome, is critical for inducing optimal pathogen-driven Th17 responses. More importantly, T cells lacking caspase-1 fail to induce colitis following transfer into RAG-deficient mice, further demonstrating the importance of caspase-1 for the development of pathogenic Th17 cells *in vivo*. This study underlines the importance of DC-mediated priming in identifying novel regulators of T cell differentiation.

## Introduction

CD4 T cells play a central role in adaptive immunity through the secretion of specific effector cytokines and also by regulating B cell activation and CD8 T cell responses (*1, 2*). Mature dendritic cells (DCs) are primarily responsible for the priming and differentiation of naïve CD4 T cells into several effector lineages. Following pathogen recognition through pattern recognition receptors (PRRs), DCs upregulate MHC Class II and costimulatory molecules and secrete innate cytokines that dictate the priming and differentiation of distinct T cell lineages (*3*). Upon receiving cues from DCs, pathogen/antigen-specific CD4 T cells rapidly proliferate and undergo transcriptional programming, including the upregulation and stabilization of the lineage-specific transcription factors (TFs, T-bet for Th1, GATA3 for Th2 and RORγt for Th17) that facilitates effector cytokine production. More specifically, Th1 cells produce IFNg which directs killing of intracellular bacteria and viruses; Th2 cells secrete type-2 cytokines like IL-4, IL-5 and IL-13 which mediate expulsion of helminths; and Th17 cells produce IL-17A, IL-17F and IL-22 which facilitate the clearance of extracellular bacteria and fungi (*4*).

The differentiation of CD4 T cells *in vivo* in response to a pathogen (here on referred to as ‘pathogen-specific T cells’) results in a heterogeneous effector population (*5*). The frequency of naïve precursors for specific epitopes is extremely low, ranging from 0.8-10 cells per million naïve CD4 T cells per epitope (*6*), making it technically challenging to detect, track and analyze pathogen-specific T cells *in vivo*. Transcriptional regulation of CD4 T cell differentiation is influenced by many factors including T cell receptor (TCR) affinity to distinct epitopes (*7, 8*). Therefore, studying transcriptional landscape of single specific TCRs may not accurately reflect the changes dictated by varying TCR strengths. Additionally, pathogen-specific CD4 T cells exhibit heterogeneity and plasticity (*9*). For example, functionally significant novel T cell subtypes (*e.g*. Th1/Th17 dual-lineage cells) are enriched in barrier tissues and can perform either pathogenic or regulatory roles (*10, 11*), but the differentiation mechanisms required to produce such dual-lineages remain largely unknown. Although DCs are major drivers of CD4 T cell activation and differentiation, lineage-specific polarization by using defined cytokine cocktails has been a major approach to study CD4 T cell biology (*4, 12, 13*). This approach fails to take into account that a broad range of DC-derived cues that might act on T cells during priming and differentiation. In addition to inducing lineage-specific TFs, DCs impact CD4 T cell differentiation by altering TCR signaling strength, licensing the expression of co-TFs or regulatory microRNA species (*8, 14*).

To understand if dendritic cells exposed to different pathogens regulate the transcriptional profile of newly primed CD4 T cells, we have utilized an *in vitro* approach to prime naive CD4 T cells. This approach allows an unbiased assessment of pathogen-directed clonal expansion and differentiation of naïve CD4 T cells. We have found that in vitro priming was able to generate pathogen-specific effector CD4 T cells and the effector lineage commitment was dictated by the nature of the priming pathogen. Comparison of the transcriptional profile of cytokine-polarized and pathogen primed Th17 cells led to the identification of a unique gene cluster associated with DC-mediated priming. We identified caspase-1 to be one of the unique genes upregulated in pathogen-primed Th17 cells but not in cytokine-polarized Th17 cells. We further established that caspase-1, independent of its role in inflammasome activation, functions in a T-cell-intrinsic fashion to promote Th17 differentiation. This study establishes that DCs provide critical cues for transcriptional programming of CD4 T cells, that are absent during cytokine-driven polarization, and furthermore provides a framework for identifying novel regulators of CD4 T cell differentiation.

## Results

### Dendritic cells exposed to pathogen lysates induce *de novo* differentiation of naïve CD4 T cells *in vitro*

To study the transcriptional regulation of pathogen-specific CD4 T cell during differentiation, we sought to design and validate an *in vitro* priming system to mimic *in vivo* priming of naïve CD4 T cells following microbial infections. We posited that *in vitro* priming would allow the identification and characterization of all pathogen-specific CD4 T cells at various stages of differentiation. In addition, we predicted that we would be able to determine the specificity of the generated responses. Dendritic cells, both migratory and lymphoid organ resident, play a central role in T cell differentiation (*3*). Therefore, a simple, isolated system that contains core elements required for T cell priming: dendritic cells, naïve CD4 T cells and complex components from pathogens, should allow the differentiation of pathogen-specific CD4 T cells. Similar co-culture approaches have been reported previously for the activation of human and mouse CD4 T cells (*15-20*). More recent work has elegantly demonstrated that exposure of human monocytes to different pathogens leads to priming of pathogen-specific CD4 T cells, whose effector cytokines are dictated by the priming pathogen (*15, 21*). However, concerns are that these studies used peripheral blood monocytes or bone-marrow derived dendritic cells (BMDCs), which are composed of a heterogeneous mixture of macrophages and ‘myeloid’ DCs with distinct antigen presentation abilities (*22-24*). These populations have been shown to deviate from lymphoid and tissue-resident DC populations (*22*). Furthermore, the pathogen specificity and functionality of the primed T cells in the previous studies was not fully established (*17-20, 25*).

We designed the *in vitro* system based on human studies using monocytes but instead using splenic CD11c^+^ DCs (Flt3l-dependent, *ex vivo*) as antigen-presenting cells (APCs) to prime naïve CD4 T cells isolated from spleen and peripheral lymph nodes (Fig. 1A) (*15*). Whole-cell lysates of bacteria including diverse PAMPs and proteins were used as PRR stimuli and source of antigens for DCs. Subsequently, DCs were co-cultured with highly purified naïve CD4 T cells to initiate their activation and differentiation (Fig. 1A and Fig. S1A; Experimental Procedures). No exogenous peptide or protein was added to ensure that TCRs would solely respond to pathogen-derived peptides. Recombinant cytokines were also not provided; thus, the effector response is not dictated by specific exogenous cytokines but solely dependent on pathogen-induced, DC-derived cytokine milieu.

**Fig. 1.**
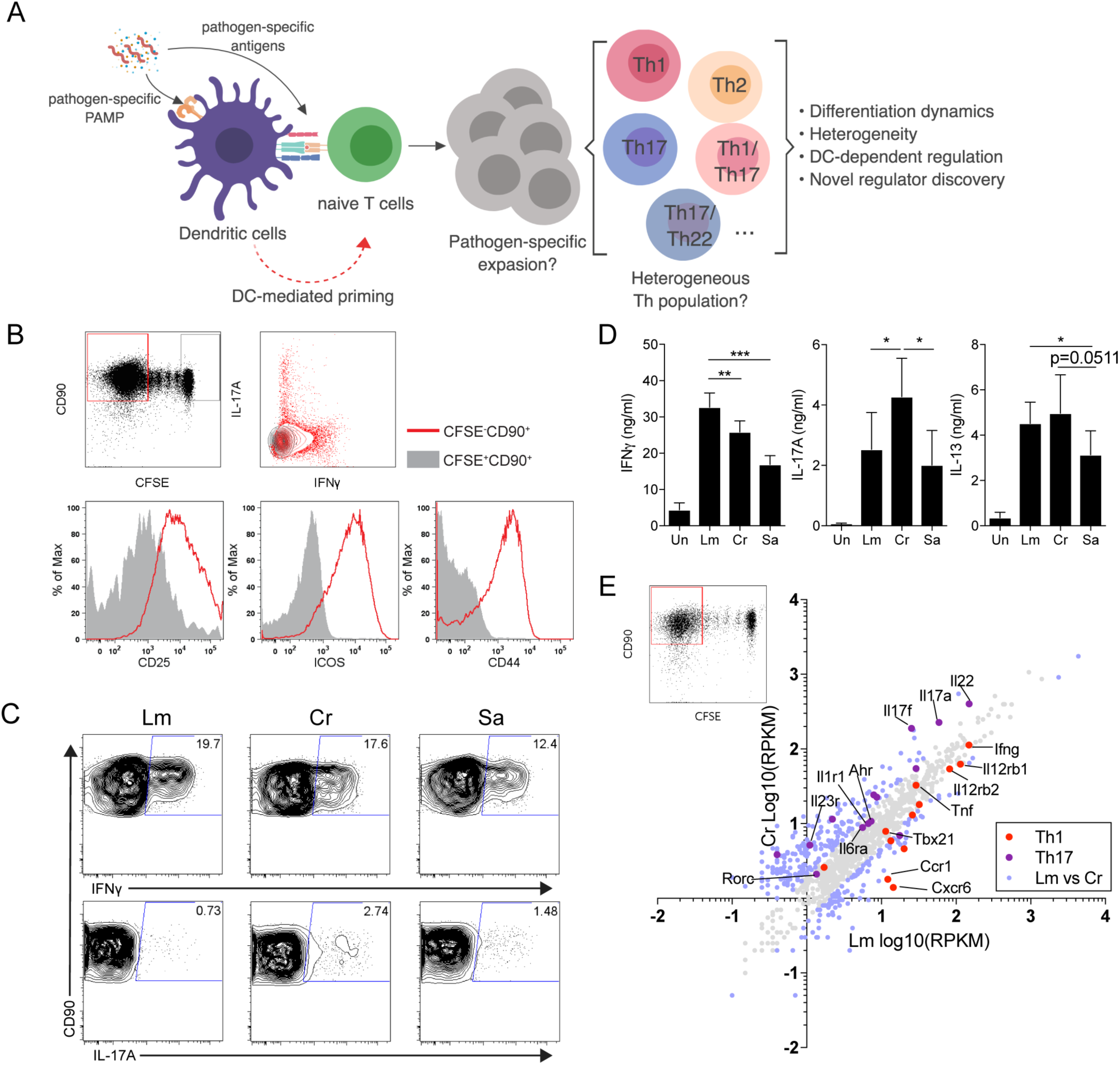
**An *in vitro* priming approach to generate functional pathogen-specific CD4 T cells** (A) Schematic overview of the priming system and workflow. (B) (upper row) Representative CFSE dilution graph and cytokine (IFNγ and IL-17A) staining from CFSE^+^ and CFSE^−^ fraction. (lower row) Histogram of CD25, CD44 and ICOS from CFSE^+^ and CFSE^−^ fraction. Cells were co-cultured for 10-12 days before analysis. (C) Representative intracellular staining of IFNγ- and IL-17A-producing CD4 T cells following priming by CD11c^+^ DCs stimulated with Lm, Cr or Sa lysates. (D) Level of IFNγ, IL-17A and IL-13 in the supernatants of co-cultures as in (C), measured by ELISA. (E) Scatter plot representing mRNA expression values (log_10_RPKM) of Lm- or Cr-specific CFSE^−^ T cells from RNA-sequencing, data are pooled and averaged from two independent samples. Blue data points indicate differentially (fold change≥2) expressed genes. All plots are pre-gated on live cells. Data are representative or pooled from at least 2 independent experiments.

Under this approach, a proportion of co-cultured naïve CD4 T cells underwent proliferation (CFSE^−^CD90^+^, Fig. 1B) and upregulated T cell activation markers CD25, CD44 and ICOS (Fig. 1B, lower). CFSE^−^CD90^+^ cells also secreted effector cytokines (Fig. 1B, upper right). T cells that failed to undergo expansion (CFSE^+^CD90^+^) remained in naïve state and did not produce effector cytokines (Fig. 1B, upper right). In contrast to non-specific polyclonal priming mediated by CD3 ligation in the presence of LPS stimulated DCs, we observed a binary distribution between CFSE^−^ cells undergoing division and static CFSE^+^ T cells (Fig. S1B), indicating clonal expansion of a small proportion of T cells. A large proportion of static CFSE^hi^ cells suggested a very low frequency of naïve T cells to have undergone active priming (Fig. S1B). Accordingly, generation of in vitro pathogen-primed T cells required pMHC-II:TCR interaction and costimulation, as well as innate cytokine milieu (for example, IL-12 for generation of IFNγ+ cells) (Fig. S1C and S1D) thus ruling out the possibility of antigen-independent, inflammatory cytokine-driven T cell proliferation (*26*).

### *In vitro* priming system generates pathogen-dictated CD4 T cell responses

To test whether this approach can induce differentiation of functional effector CD4 T cells *in vitro*, we took advantage of three well-studied mouse pathogens: *Listeria monocytogenes* (Lm), *Citrobacter rodentium* (Cr), and *Staphylococcus aureus* (Sa). The level of proliferation of CD4 T cells primed by different pathogen lysates was comparable (Fig. S1E). However, CFSE^−^ CD4 T cells exhibited distinct cytokine profiles: Cr-primed T cells generated significantly more IL-17A^+^ population compared to both Lm and Sa; Lm-priming generated predominantly IFNγ-producing CD4 T cells and Sa-stimulated DCs primed both IFNγ^+^ and IL-17A^+^ CD4 T cells (Fig. 1C). This was also evident when we measured these cytokines in the supernatants (Fig. 1D). Th1/Th17 profile observed in the *in vitro* system are in concordance with the dominant responses previously reported for each pathogen *in vivo* (*27-29*). Live infection of DCs with these pathogens also led to similar Th1/Th17 profile as lysates and therefore we used bacterial lysates for rest of the study to be consistent for antigen dosage and concentration or PRR ligands (Fig. S1C).

We sorted the total CFSE^−^ CD4 T cell population from Lm- and Cr-primed cultures and performed mRNA sequencing to compare their global transcriptional profiles (Fig. 1E). Cr-primed T cells revealed a Th17-associated gene signature, while Lm-primed T cells predominantly contained a high level of Th1-associated transcripts. Notably, 126 genes were uniquely expressed in Lm-primed T cells and 233 genes were uniquely expressed in Cr-primed T cells, when normalized to their CFSE^+^ naive counterparts (Fig. 1E; Table S3), indicating pathogen-associated T cell transcriptome. These experiments established that dendritic cells, exposed to a complex mixture of pathogen proteins and PRR ligands, can indeed prime naïve CD4 T cells *in vitro*. More importantly, depending on the nature of the pathogenic stimuli, DCs were able to shape distinct CD4 T cell transcriptional profiles.

### *In vitro* pathogen-primed T cells are specific to the priming pathogen

To interrogate whether these in vitro primed T cells were specific to the priming pathogen, we first examined TCR repertoire enrichment by comparing CFSE^−^ primed cells to CFSE^+^ naïve pool (Fig. S2A). We observed a selective enrichment of TCR Vβ10,12,14 and 15 in CFSE^−^ population. Meanwhile, expression of Vβ13-3, 20, 26 was diluted in CFSE^−^ population, indicating a lack of proliferation of these Vβ-expressing T cells in response to Lm (Fig. S2A). To measure specificity, we rested pathogen-primed CD4 T cells and then restimulated these cells *in vitro* with irradiated, pathogen lysate fed B cells. We found that primed and rested CD4 T cells secreted IFNγ and IL-17A only when re-stimulated by B cells exposed to the original priming pathogen (Fig. 2A and S2B). Cytokine production and upregulation of T cell activation markers (CD25, CD69, ICOS and CD44) were dependent on the dose of lysates used for reactivation, further confirming the specificity of the responding T cells (Fig. 2A-B and S2B-C). In contrast, restimulation with the non-priming (mismatched) pathogen did not lead to cytokine secretion at any of the doses used (Fig. 2A-B and S2B-C), further establishing the specificity of the response.

**Fig. 2.**
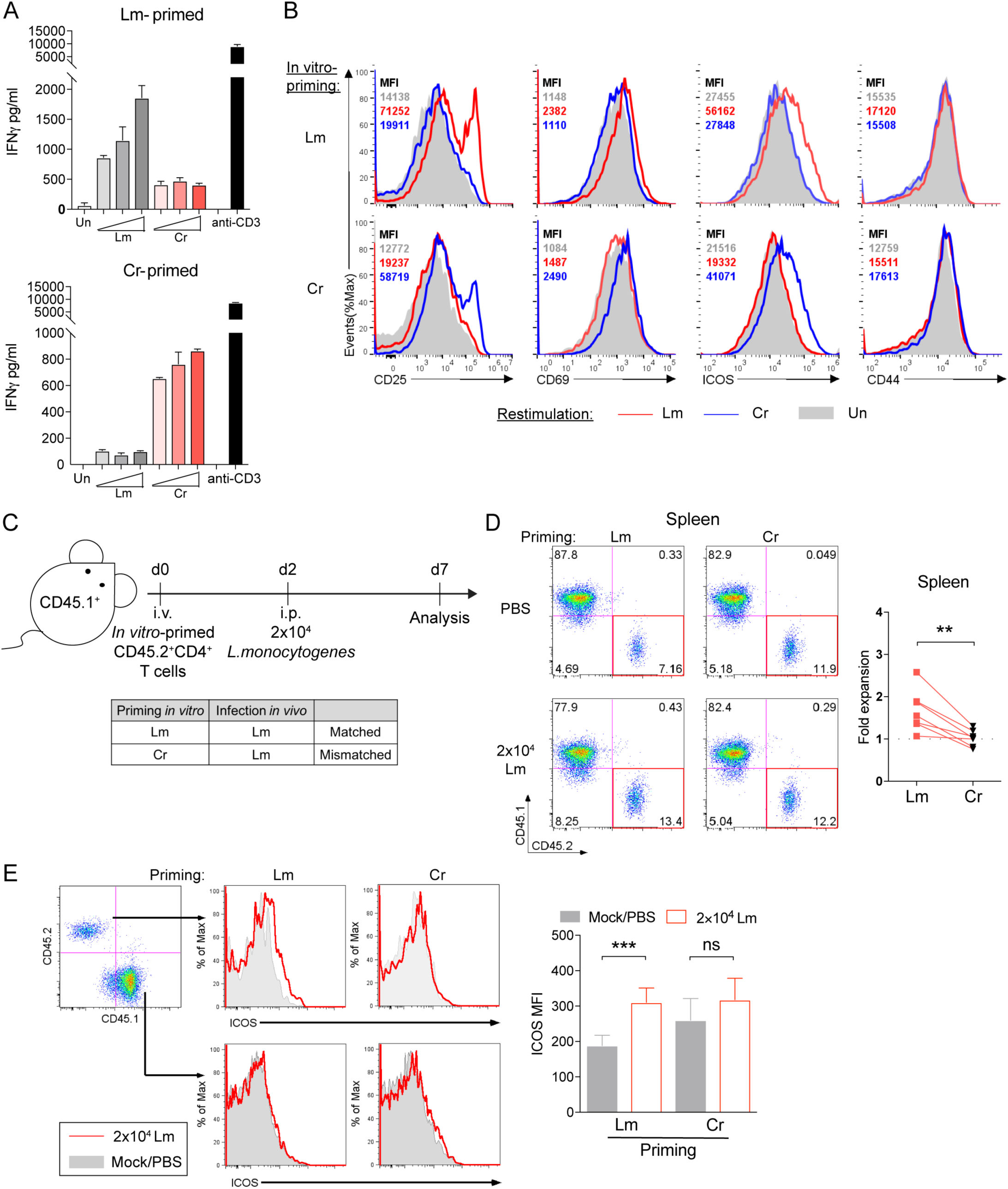
***In vitro* pathogen-primed CD4 T cells exhibit specificity towards the priming pathogen** (A) IFNγ quantities in the culture supernatant of Lm- (upper panel) or Cr- (lower panel) primed CD4 T cells that were cultured for 48 hours with unstimulated or Lm/Cr-fed, irradiated B cells. Lm and Cr concentrations used for restimulation were titrated at 3, 10 and 30µg/ml. Data are representative of 4 independent experiments. Culture supernatants from anti-CD3 (30 ng/ml) stimulated T cells we also assessed for IFNγ production as the positive control. Data are representative of 3 independent experiments. (B) Histogram and MFI (upper left corner) of CD25, CD69, ICOS and CD44 on the CD90^+^ T cells from the same experiments in Figure 2A denoting upregulation of indicated activation markers in response to Lm/Cr rechallenge. Lysate concentration=10µg/ml. Data are representative of 2 independent experiments. (C) (upper) Experimental design for testing *in vivo* specificity and (lower) mismatch scheme. (D) Representative flow cytometry analysis and quantified percentages (right) of transferred (*in vitro* primed with Lm or Cr) CD4 T cells in the spleen at day 5 post-infection (shown as fold change comparing CD45.2^+^% of infected mouse to paired PBS control). n=7 mice per group. (E) Mean fluorescent intensity (MFI) of surface ICOS on donor CD45.2^+^ T cells and recipient endogenous CD45.1^+^ T cells from the same experiments in Figure 2D and quantified for CD45.2^+^ T cells (right). n=7 mice per group. Error bars represent mean ± SEM and p values were determined by paired Student’s t-test. ^∗∗^p < 0.01, ^∗∗∗^p < 0.001.

Using a similar mismatch scheme (Fig. 2C), we further tested whether *in vitro* primed CD4 T cells respond to infection by the priming pathogen *in vivo*. To that end, we transferred Lm- or Cr- primed CD45.2^+^ T cells to CD45.1^+^ congenic, immune competent recipients and recipients were challenged with intraperitoneal Lm infection 48 hours after transfer (Fig. 2C and S2D). Mice that received pathogen-primed CD4 T cells but were not subsequently infected with *L.monocytogenes* served as baseline controls. Donor CD4 T cells proliferated in response to matched pathogen re-challenge (Lm-primed donor cells and Lm infection; Fig. 2C) but did not respond to mismatched pathogen re-challenge (Cr-primed donor cells and Lm infection) in the spleen (Fig. 2D) and lymph nodes (Fig. S2E). In addition, matched re-challenge induced significant upregulation of cell surface ICOS expression on donor cells (Fig. 2E). ICOS upregulation on recipient CD45.1^+^ CD4 T cells was unaffected by the transfer of Lm- or Cr- primed T cells (Fig. 2E and S2F). It is important to note here that we used a very low dose of Lm for challenge and responses were assessed shortly after infection that precluded the possibility of *de novo* priming of naïve CD4 T cells in the recipient. In summary, these data allow us to conclude that the *in vitro* priming system generates effector T cells bearing TCRs that are likely to be reactive to peptides derived from the original priming pathogen.

## Pathogen-primed Th17 cells are comprised of heterogeneous subsets and display a transcriptional profile different from cytokine-polarized Th17 cells

Among all the effector T cell lineages, Th17 cells have been reported to have a high degree of plasticity and heterogeneity (*30*). They are sensitive to variations in polarizing cytokines between distinct microenvironments (*31*), and can convert to regulatory subsets during the resolution of inflammation (*32*). This intrinsic fine-tuning of Th17 cells makes them an ideal effector lineage to dissect their transcriptional regulation during pathogen-induced differentiation. We generated mice that would allow fate-mapping of CD4 T cells that commit to making IL-17A (‘17A-fm’ mice; *Il17a-cre*^+/−^; *Rosa26-flox-stop-flox-tdTomato*^+/−^) (*33*). Cr-primed CD4 T cells contained a higher proportion of tdTomato^+^ cells than Lm-primed CD4 T cells (Fig. S3A), consistent with our previous data (Fig. 1C), indicating successful tracing of Th17 cells committed for IL-17A production.

Traditional cytokine-differentiated Th17 cells (cdTh17) have been widely used for global transcriptional profiling (*12, 13*). To understand transcriptional programming of pathogen-specific Th17 lineage cells, we used Cr-primed DCs to differentiate 17A-fm naïve T cells or polarized 17A-fm naïve T cells to Th17 lineage using antibodies and cytokine cocktails (Fig. 3A). tdTomato^+^ populations, FACS-sorted from both conditions, were subjected to RNA-sequencing analysis. Clustering of Th17 associated cytokine and TF expression confirmed that tdTomato^+^ population in pathogen-primed cultures are highly enriched for Th17 cells, compared to CFSE^+^ or CFSE^+^ tdTomato^−^ populations (Fig. S3B). Principal component analysis (PCA) demonstrated a distinct transcriptional profile between cytokine-differentiated Th17 cells and Cr-primed Th17 cells (Fig. 3B and Table S4). Pathway analysis revealed that ppTh17 cells expressed high levels of genes related to diverse T cell effector functions while cdTh17 cells upregulated cell cycle- and proliferation-related genes (Fig. 3C and Table S5). Consistently, cdTh17 cells exhibited a strongly activated phenotype characterized by the extremely high expression of Th17-lineage master TFs *Rorc, Rora* and *Ahr* and the enhanced expression of Th17 lineage cytokines *Il17a, II17f* and *Il22* (Fig. 3D). In addition, cdTh17 cells significantly upregulated *Il9*, known to further promote Th17 lineage (*34*) (Fig. 3D). In contrast, ppTh17 cells exhibited a multi-lineage phenotype by expressing an array of Th1 and Th2-associated genes, such as transcription factor T-bet (*Tbx21*), GATA3 (*Gata3*) and cytokines (*IFN_γ_, II4, II5, II13*), representing potential lineage plasticity (*35*) (Fig. 3D). Interestingly, cdTh17 cells expressed a high level of transcription factor *Foxp3*, marking TGFβ induced co-differentiation of regulatory T cell lineage with Th17 lineage (*36*). In comparison, ppTh17 expressed a decreased level of *Foxp3*, indicating that Cr-specific T cells do not transdifferentiate to Tregs. To test whether Cr-specific Th17 cells transdifferentiate into Th1 cells, we primed naïve T cells from 17-γ double reporter mice (*Il17a-cre*^+/−^; *Rosa-flox-stop-flox-tdTomato*^+/−^; *Ifng-ires-yfp*^+/−^) with Cr. From day 5 post stimulation, we observed the emergence of tdTomato^+^YFP^+^ cells indicating emergence Th1 transdifferentiated cells from T cells previously committed to Th17 lineage (Fig. 3E). The proportion of this population increased as the cells proliferated over an extended period of differentiation (from day 5 to day 10) (Fig. 3E). In contrast, cdTh17 cells did not re-express YFP, even after the removal of polarizing cytokines (Fig. 3E). Although studies have shown polarized Th17 cells are able to transit to Th1 cells upon secondary stimulation (*37, 38*), our data indicate that plasticity during primary commitment is limited in cdTh17 cells.

**Fig. 3.**
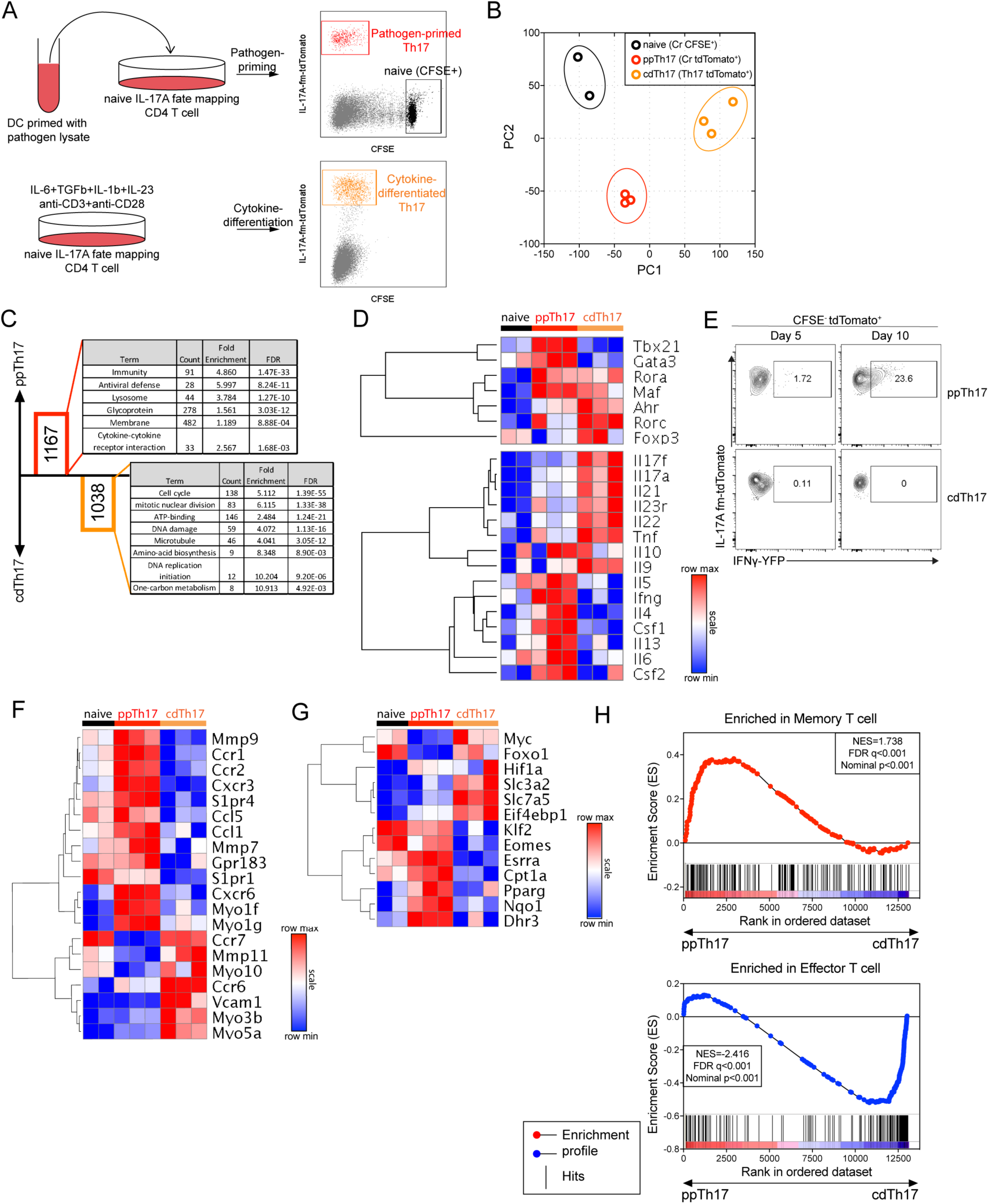
**Comparative transcriptional analysis reveals major divergence in programming between pathogen-primed Th17 and cytokine-polarized Th17 cells** (A) Experimental design for transcriptome profiling of pathogen-primed or cytokine-differentiated Th17 cells. (B) PCA analysis of whole transcriptome expression of CFSE^+^ (naïve), Cr-specific (ppTH17, day 12) or cytokine-differentiated (cdTh17) Th17 cells. Each data point indicates one independent replicate. (C) The number of genes specifically upregulated in ppTh17 (red) or cdTh17 (orange) cells and their functional annotation enrichment analyzed by DAVID. (D) Heatmap and hierarchical analysis of key T cell transcription factors, cytokines and cytokine receptor expression from transcriptome profiling described in Figure 3A. Replicates are shown in each column. (E) Representative flow plots showing YFP^+^% of CFSE^−^tdTomato^+^ population, from 17-γ double reporter T cells under ppTh17 (Cr-primed) or cdTh17 conditions at early (day 5) and late (day 10) stage of differentiation. For day 10 cdTh17 cells, polarizing cytokines were removed from the culture after day 5 and cells are maintained in 10ng/ml rIL-2 media from day 5 to day 10. Data are representative of two independent experiments. (F) Heatmap and hierarchical analysis of gene expression for gene cluster involved in *in vivo* T cell motility, migration, chemokine and chemokine receptor signaling, T cell positioning and antigen sampling. (G) Heatmap and hierarchical analysis of gene expression for genes involved in metabolic processes. (H) GSEA analysis of ppTh17 and cdTh17 cells compared to Molecular Signature dataset of effector versus memory T cells. Hierarchical clustering was determined by Pearson correlation and pairwise average-linkage.

Further analysis of the transcriptome revealed that ppTh17 cells upregulated a set of genes encoding membrane-associated proteins that are important for chemotaxis (Fig. 3F, Table S5), including chemokine receptors (*Ccr1* and *Cxcr3*), extracellular matrix metalloproteases (*Mmp7* and *Mmp9*), myosins (*Myo1f* and *Myo1g*), S1P receptor family (*S1pr1* and *S1pr4*) and G-protein-coupled receptor EBI2 (*Gpr183*) (Fig. 3F and Table S6) (*39-42*). Interestingly, we also found highly enriched interferon-stimulated genes (ISGs) in naïve and ppTh17s (Fig. S3C), as IFI16 (*Ifi204*) and STING-activation has been demonstrated to mediate anti-proliferative effect in CD4 T cells and promote memory formation (*43*). We sought to determine whether transcripts of ppTh17s denote the acquisition of memory T cell state. Comparison of metabolic gene datasets indicates that ppTh17 cells upregulated AMPK pathway, fatty acid oxidation and oxidative phosphorylation programs that maintain memory T cell function (Fig. 3G) (*44, 45*); cdTh17 cells upregulated c-Myc and HIF1α targets and resemble terminally differentiated effector T cells that employ glycolysis as energy fuel (Fig. 3G) (*46-48*). Consistently, ppTh17 expressed genes upregulated in memory T cells, such as *Il7r and Il15ra* (Fig. S3D), which are important for the maintenance of memory T cells through IL-7R and IL-15R signaling (*49, 50*). To further test our hypothesis on a global transcriptional landscape, we performed gene set enrichment analysis (GSEA) with memory versus effector T cell molecular signature from ImmuneSigDB database (*51*). GSEA analysis indicated that memory T cell-associated genes were enriched in ppTh17s compared to cdTh17s (Fig. 3H, upper). Effector T cell-associated genes were enriched in cdTh17s, consistent with their metabolic status (Fig. 3G and H, lower; Table S7).

We further validated some highly upregulated genes in ppTh17 compared to cdTh17 by quantitative RT-PCR (Fig. S3E). Since the experiments so far have focused on identifying distinct transcriptional profiles between pathogen-primed or cytokine-polarized CD4 T cells *in vitro*, we also assessed the physiological relevance of our findings. We infected the 17A-fm reporter mice with *C. rodentium*, sorted total CD4^+^tdTomato^+^ cells (consist of differentiated, Cr-specific Th17 cells) from mesenteric lymph nodes (mLNs) at the peak of infection, and assessed the expression of these key variable genes (Fig. S3E and S3F). The majority of the genes tested had comparable expressions between CD4^+^tdTomato^+^ cells and ppTh17 cells. Furthermore, unsupervised hierarchical clustering showed that ppTh17s resembled *in vivo* effector Th17 cells (Fig. S3F), indicating a close functional relationship between ppTh17 cells and *in vivo* generated Th17 cells during infection.

In summary, we find that in contrast to cytokine-polarized Th17 lineage cells that display features of terminal differentiation, DC-mediated priming led to Th17 lineage cells of higher plasticity and heterogeneity as observed in *in vivo* primed T cells.

## Identification of caspase-1 as a DC-induced, T-cell-intrinsic regulator of pathogen-specific Th17 cell differentiation

Our analysis (Fig. 3B-D) and some other evidence suggest DCs directly modulate the heterogeneity and plasticity of Th17 cells (*15*). Whether DCs regulate Th17 responses through the expression of specialized regulators is still unclear (*52*). To identify such regulators, we focused on genes that would be uniquely expressed in ppTh17 but not in cdTh17 cells. Surprisingly, *Casp1* (encoding caspase-1) emerged as an interesting candidate, with 5- to 10-fold induction of expression in ppTh17 cells compared to cdTh17 cells or naïve T cells (Fig. 4A and S3E). We also found that *Casp1* is upregulated in *ex vivo* effector T cell population primed in mLNs following *C. rodentium* infection (Fig. S3F). Caspase-1 and caspase-11 (gene name as *Casp4*) are inflammatory caspases that have overlapping effector functions downstream of inflammasome activation primarily in myeloid cells (*53, 54*). However, the role of these caspases in T cells is not well defined. *Casp1* and *Casp4* transcripts are upregulated in memory T cells compared to naïve T cells in Immgen database (www.immgen.org; Fig. S4A). However, *Casp4* levels are not significantly different between ppTh17 and cdTh17 (Fig. 4A and S4B), indicating that *Casp4* may be upregulated upon TCR activation but *Casp1* is induced in T cells by unknown cues from DCs.

**Fig. 4.**
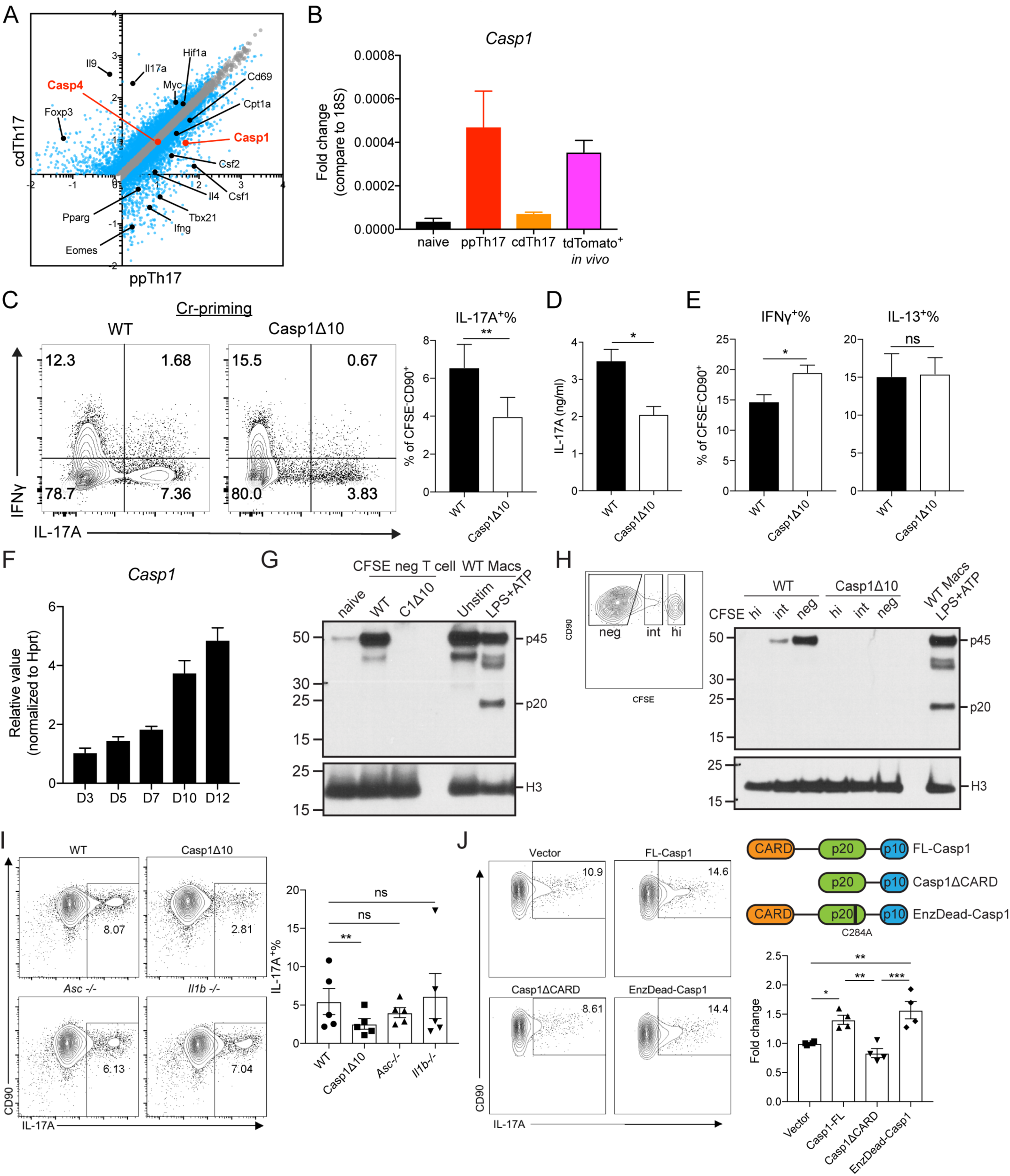
Caspase-1 promotes the differentiation of Th17 lineage independent of its enzymatic activity or inflammasome activation. (A) Differentially expressed transcripts between ppTh17 and cdTh17 cells. Each dot represents the average of three independent experiments. Blue dots indicate differentially regulated genes (fold change>1.5, FDR<0.05). Black dots indicate differentially regulated transcripts described in Figure 3. Red dots indicate Casp1 and Casp4 transcripts. (B) mRNA Expression of *Casp1* in sorted naïve (CFSE^+^), ppTh17, cdTh17, or tdTomato^+^ cells from mLNs of Cr-infected (day-10 p.i.) 17A-fm mice, quantified by independent qPCR experiments. n=2. (C) Naïve CD4 T cells from WT or Casp1Δ10 mice were primed *in vitro* by Cr lysate-stimulated WT splenic CD11c^+^ dendritic cells, IL-17A and IFNγ productions were measured by intracellular staining and flow cytometry analysis of CD90^+^CFSE^−^ live CD4 T cells (C, left); IL-17A^+^% are quantified (C, right). n=7 experiments. (D) IL-17A in the supernatant from experiments in (C), measured by ELISA. n=3. (E) IFNγ^+^% and IL-13^+^% of CD90^+^CFSE^−^ live cells, quantified from experiments in (C). n=7. (F) Relative expression of *Casp1* mRNA at indicated time points post Cr-priming. (G) Western blot analysis of pro-caspase-1 (p45) and cleaved caspase-1 (p20) from naïve CD4 T cells, sorted CD90^+^CFSE^−^ Cr-primed CD4 T cells (day-12), bone marrow-derived macrophages that were unstimulated or under conventional inflammasome activation (4hr LPS+30min ATP). (H) Western blot analysis of pro-caspase-1 (p45) and cleaved caspase-1 (p20) of Cr-primed CD4 T cells. Cells are sorted into CFSE^high^, CFSE^intermediate^ and CFSE^low^ populations (left) and compared to WT macrophages undergoing inflammasome activation. (I) Intracellular cytokine staining of WT, Casp1Δ10, *Asc*^−/−^, *II1b*^−/−^ CD4 T cells, primed with Cr-stimulated DCs (left). IL-17A^+^% of CFSE^−^CD90^+^ live cells was quantified (right). n=5. (J) Casp1Δ10 CD4 T cells were differentiated to Th17 lineage and retrovirally reconstituted with MSCV-IRES-hCD2 alone (Vector), full-length Casp1 (FL-Casp1), Casp1 deficient of CARD (Casp1ΔCARD) or enzymatically inactive form of Casp1 (EnzDead-Casp1, C284A) and quantified for IL-17A^+^% (gated on live, hCD2^+^ population). n=4. Statistics represent mean ± SEM and p values were determined by paired Student’s t-test. ^∗^p<0.05, ^∗∗^ p < 0.01, ^∗∗∗^p < 0.001.

To test whether DC-induced caspase-1 plays a role in pathogen-specific Th17 differentiation, we used Casp1Δ10 mice which specifically lack the expression of caspase-1 but not caspase-11 (*55*). We found Casp1Δ10 T cells to be defective in Th17 differentiation following *in vitro* priming with Cr-stimulated WT DCs (Fig. 4C and D). Commitment to Th1 or Th2 lineages in Casp1Δ10 CD4 T cells was unaffected (Fig. 4E). There was no difference in proliferation between WT and Casp1Δ10 T cells during *in vitro* priming (Fig. S4C). These results suggest a highly critical role for T-cell-intrinsic caspase-1 in inducing optimal Th17 response during pathogen-mediated differentiation.

## Caspase-1 functions in CD4 T cells independently of its canonical enzymatic activity or inflammasome activation

The role of caspase-1 in inducing pyroptotic cell death in HIV-infected CD4 T cells has been previously demonstrated (*56, 57*). However, the role of T-cell-intrinsic caspase-1 in normal CD4 T cell differentiation is still unclear. When we examined the expression pattern of *Casp1* transcripts during differentiation, we found that *Casp1* mRNA increased temporally, correlating with the increase of IL-17A-fm-tdTomato signal (Fig. 4F and S4D). Similarly, we found caspase-1 protein to be expressed highly after day 7 post stimulation and is maintained through day 12 with the highest levels expressed in CFSE^−^ differentiated population (Fig. 4G-H and S4E). However, in primed T cells, we did not find cleaved caspase-1 p20, an active product resulted from inflammasome activation in macrophages (Fig. 4G-H). The absence of cleaved form suggests that the role of caspase-1 in CD4 T cells could be independent of its canonical function in the inflammasome complex as an IL-1 cleaving enzyme (*58*). Consistent with this idea, we did not observe a defect in Cr-induced priming in *Asc*^−/−^ and *II1b*^−/−^ CD4 T cells (Fig. 4I). IL-1β, secreted following inflammasome activation, has been shown to promote Th17 responses *in vivo* (*59*). However, supplementing IL-1α or IL-1β to Casp1Δ10 T cells cultures failed to rescue the defect in IL-17A^+^ cell differentiation (Fig. S4F), suggesting a function for caspase-1 that is independent of its IL-1β cleavage activity.

Given that we did not observe cleaved caspase-1 in CD4 T cells, we looked into a possibly catalytically-independent function of caspase-1. Caspase activation and recruitment domain (CARD) of caspase-1 facilitates homeostatic binding to effector proteins such as ASC and RIP2 (*54*). Binding of caspase-1 to RIP2 through CARD, for example, promotes NF-kB signaling independent of its enzymatic activity (*60*). To test the possible scaffolding function of caspase-1, we ectopically reconstituted Casp1Δ10 CD4 T cells with full-length (FL), CARD-deficient (ΔCARD) or enzymatically inactive (EnzDead) caspase-1 and investigated their ability to induce Th17 lineage commitment (Fig. S4G). Reconstitution of FL and EnzDead caspase-1 resulted in an increase of IL-17A-producing T cells compared to vector alone but ΔCARD failed to promote Th17 differentiation (Fig. 4J). EnzDead caspase-1 led to modestly enhanced Th17 differentiation than FL-caspase-1, suggesting that inhibiting catalytic activity might promote functions associated with the CARD domain of caspase-1 (Fig. 4J). Overall these data provide compelling evidence that T-cell-intrinsic caspase-1, in an inflammasome-independent fashion, promotes Th17 differentiation.

## T-cell-intrinsic caspase-1 is required for Th17-mediated disease *in vivo*

Even though we found that absence of T-cell-intrinsic caspase-1 affected the generation of pathogen primed Th17 cells, TCR activation- and cytokine cocktail-driven Th17 polarization (Fig. S5A) and proliferation (Fig. S5B) was unaffected. This prompted us to examine the *in vivo* relevance of our findings. By analyzing steady-state T helper cell populations in co-housed WT and Casp1Δ10 mice, we found reduced IL-17A^+^% (Fig. S5C) and IL-22^+^% (Fig. S5D) CD4 T cells in the spleen, mLN and small intestine lamina propria (LP) of Casp1Δ10 mice, consistent with previous reports (*61*). The percentage of IFNγ-producing CD4 T cells was unchanged in mLN and LP but increased in the spleens of Casp1Δ10 mice (Fig. S5E). Since these outcomes could be a result of caspase-1 deficiency in myeloid cells, we further investigated the role of T cell autonomous caspase-1 *in vivo* by using a T cell transfer model of colitis. In this approach, highly purified naïve CD4 T cells from WT and Casp1Δ10 mice are transferred to *Rag1*^−/−^ recipients thus restricting the genetic deficiency to the CD4 T cell compartment. Since transferred naïve CD4 T cells differentiate in response to components of gut microbiota and germ-free mice develop a very mild disease (*62, 63*), this approach would allow us to test the significance of our *in vitro* priming system.

We found that *Rag1*^−/−^ animals that received Casp1Δ10 naïve CD45RB^hi^ CD4 T developed very mild disease when compared to WT T cell recipient mice that developed measurable colitis, including significant weight loss (Fig. 5A) and disease progression (Fig. 5B). Recipients of WT T cells also showed significant colon shortening compared to non-T cell transferred controls (Fig. 5C). However, transfer of Casp1Δ10 T cells did not lead to this disease manifestation (Fig. 5C). Transfer of WT T cells also led to severe colonic pathology, marked by transmural infiltration of leukocytes, epithelial cell hyperplasia and submucosal morphological changes, while transfer of Casp1Δ10 naïve T cells induced significantly less leukocyte infiltration and morphological changes associated with colitis (Fig. 5D and S5F). We observed significantly lower IL-17A^+^IFNγ^+^ in mLNs and colons of Casp1∆10 T cell recipients compared to WT T cell recipients (Fig. 5E). These IL-17A^+^IFNγ^+^ pathogenic Th17 cells have been demonstrated to be critical for the induction of T cell mediated-colitis and development of pathology (*10*). Of note, there was no difference in the proportion of IL-17A^+^IFNγ^−^ non-pathogenic population between WT and Casp1Δ10 recipients (Fig. S5G). In contrast, we observed the significantly less splenic expansion of total IL-17A^+^ Casp1Δ10 cells (Fig. 5F), consistent with the reduced circulating IL-17A levels throughout the course of the disease (Fig. 5G). Overall these data support that T-cell-intrinsic caspase-1 selectively controls the differentiation of both pathogen-specific and auto-inflammatory Th17 lineage cells. Furthermore, even though cells of both innate and adaptive immune system express caspase-1, these data provide compelling evidence for a distinct role for caspase-1 in regulating Th17 biology.

**Fig. 5.**
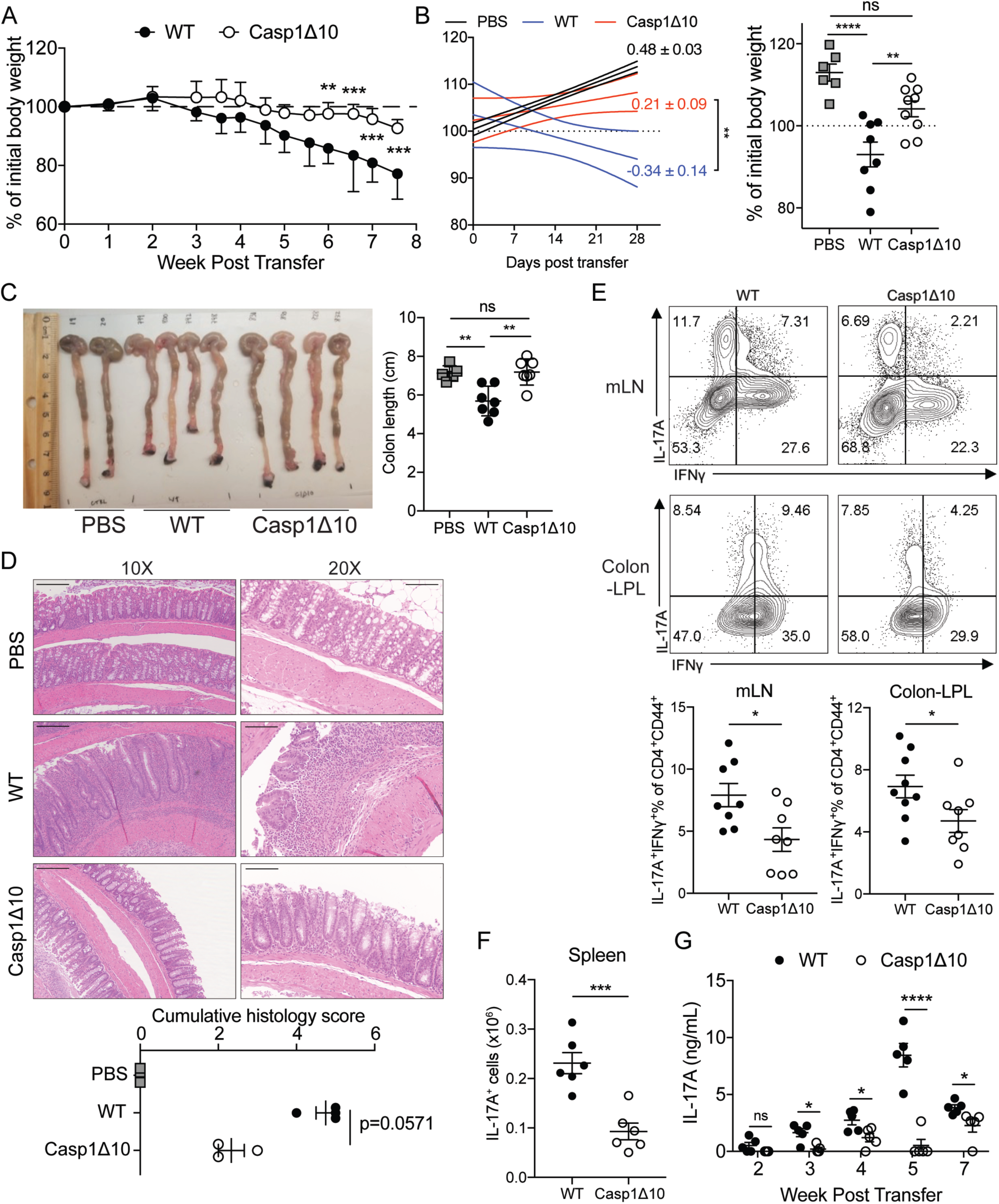
**T-cell-intrinsic caspase-1 is required for Th17-mediated colitis *in vivo*.** (A) Weight change of *Rag1*^−/−^ mouse that received WT or Casp1Δ10 naïve CD4 T cells (CD45RB^hi^) at indicated time points (n=5 mice for each genotype). (B) (left) Linear regression analysis of weight loss progression 0-4 weeks post cell transfer, progression slope ± SE was shown on the side of each curve. (right) percentage of initial body weight after 4 weeks. PBS n=6, WT n=8, Casp1Δ10 n=9. (C) Representative picture of colons (left) and measured colon length (right) of *Rag1*^−/−^ mice at 8-weeks post naïve CD4 T cell (CD45RB^hi^) transfer (PBS, n=5; WT, n=7; Casp1Δ10, n=7). (D) Representative ΗΕ staining of colon sections from PBS, WT or Casp1Δ10 naïve T cell transferred *Rag1*^−/−^ mice. Images are displayed at 10x magnification to show multiple colonic regions and 20x magnification to show one consecutive section. Histology slides were blind-scored by a pathologist at UT Southwestern (lower panel). PBS, n=5; WT, n=4; Casp1Δ10, n=3. (E) Representative flow plots showing the percentages of IL-17A^+^IFNγ^+^ of CD4^+^CD90^+^CD44^+^ T cells in the mesenteric lymph nodes (mLN) or colonic lamina propria (Colon-LPL) of *Rag1*^−/−^ mice 4 weeks post transfer of WT or Casp1Δ10 naïve CD4 T cells. mLN: WT or Casp1Δ10 n=8. Colon-LPL: WT n=9; Casp1Δ10 n=8. (F) The number of CD4^+^CD90^+^IL-17A^+^ T cells in the spleens of the *Rag1*^−/−^ mice that received WT or Casp1Δ10 naïve CD4 T cells. n=6 per group. (G) Serum IL-17A levels at the indicated time points from the mice in Figure 6F. n=5 mice per group. Data are representative of 2-3 independent experiments. Statistics represent mean ± SEM and p values were determined by two-way repeated ANOVA with Bonferroni correction (A), unpaired Student’s t-test (B, D-F) or Mann-Whitney U test (C) ^∗^p<0.05, ^∗∗^ p < 0.01, ^∗∗∗^ p < 0.001, ^∗∗∗∗^p<0.0001.

## Discussion

Although dominant and protective CD4 T cell responses to specific pathogens are well understood, transcriptional profiling of newly differentiated pathogen-specific CD4 T cells, following *in vivo* infections, has been lacking due to the absence of tools to identify all responding CD4 T cells. A recent method has highlighted an *in vitro* priming and *de novo* differentiation of naïve pathogen-specific human CD4 T cells by autologous pathogen-stimulated monocytes (*15, 21*). This isolated priming system is freed from the use of exogenous polarizing cytokines and strong or repeated antigenic exposure, allowing mapping of a primary, pathogen-specific CD4 T cell response. Combining the *in vitro* priming system with genetic fate-mapping and RNA-sequencing, we were able to track and profile the differentiation of murine pathogen-specific CD4 T cells. Consistent with previous studies, we found that while CD4 T cells primed against a particular pathogen differentiate into one dominant effector lineage, other subtypes also co-exist (Fig. 1C-E) (*5, 15*). In addition, transcriptional analysis of fate-mapped Th17 cells suggests the existence of multiple sub-lineages (Th2/Th17, IL-10- or GM-CSF- secreting populations) (Fig. 3D). The heterogeneity of the T cell response is presumably dictated by DCs encountering a complex mixture of PAMPs unique to each pathogen (*3*). But why and how only certain bacteria species induce Th17 cells still remains unclear. In addition to *C.rodentium*, several other mucosal pathogens (*e.g*. EHEC) and commensals (*e.g*. Segmented Filamentous Bacteria) favor a Th17-dominant response, and this ability was attributed to their interaction with distinct intestinal microenvironments (*31, 64-67*). Interestingly, we observed that both live Cr- and Cr lysate-stimulated splenic CD11c^+^ DCs are sufficient to induce Th17 cells (Fig. 1C and S1F), suggesting that in addition to the unique mucosal microenvironment, evolutionarily conserved ligands in certain gut-associated bacteria could promote a Th17 response. The molecular mechanisms that drive this tailored response require further investigation.

Our data suggest that DCs influence transcriptional programming of T cells that extends beyond activation and effector cytokine production. CD4 T cells differentiated by pathogen-stimulated DCs showed transcriptional profiles that display lineage plasticity and features of memory differentiation (Fig. 3). DC-dependent programming also promotes CD4 functional heterogeneity and expression of genes associated with of T cell migration and motility (Fig. 3). In line with our idea that DC-induced transcriptome contains context-specific genes that fine-tune CD4 T cell response, we found induction of caspase-1 only in T cells that were primed by pathogen-stimulated DCs. Our data also firmly establish that caspase-1, independent of its role in inflammasome activation and IL-1β production, plays a T-cell-intrinsic role for the differentiation of Th17 lineage cells. Perplexingly, cytokine-mediated polarization of Th17 cells neither induce caspase-1 nor need caspase-1 for optimal differentiation, thus highlighting the fact that these cells might not resemble physiologically generated Th17 cells. The results from the T cell transfer model of colitis extended the role of caspase-1 to the differentiation of auto-inflammatory Th17 cells. Our data also redefine the role of caspase-1 in Th17-mediated inflammatory disease *in vivo*. Earlier studies have reported that caspase-1 knockout animals have ameliorated disease in several chronic autoimmune models (*68-70*). At the time this was attributed to abrogated IL-1β and IL-18 production during inflammasome activation in myeloid cells (*69, 71*). Our results suggest the possibility that T-cell-intrinsic deficiency of caspase-1 impairing Th17 commitment could have also contributed to the phenotypes. Since the majority of the studies on inflammasome components harnessed whole body deficient mice, further efforts would be necessary to dissect the specific roles of caspase-1 in innate and adaptive immune compartments in regulating T cell differentiation and Th17-associated disease outcomes.

Despite the report that the effector function of human Th1 cells is enhanced by complement-driven, caspase-1-dependent inflammasome activation within CD4 T cells (*72*), we observed a moderately increased Th1 commitment in Casp1Δ10 T cells in mouse (Fig. 4E and S5E). Importantly, we demonstrated that the role of caspase-1 in controlling Th17 differentiation is independent of its enzymatic activity and depends on its CARD domain. Recent work has shown that caspase-1 is expressed but dispensable for IL-1β production in Th17 cells (*73*), as we found no difference in the ability of full-length or enzymatically inactive caspase-1 in driving Th17 differentiation. Additionally, caspase-1 seems to have a critical function for regulating Th17 differentiation by possibly interacting with other proteins in the cytosol using its CARD domain. A variety of proteins have been reported to contain core CARD domains, such as BCL10 and AIRE (*74, 75*), posing a wide possibility for the interaction partners for caspase-1. The putative interaction partner itself may also be regulated by an appropriate differentiation signal, in order for caspase-1 to function only in DC-mediated Th17 priming. Evolutionarily, caspase-1 in early vertebrates lacks the ability to cleave IL-1 (*76*), suggesting a possible regulatory trait of these proteins. Indeed, other CARD-containing proteins have important roles beyond the involvement in inflammatory and apoptotic cell death, particularly in linking T cell receptor activation to signaling cascade (*77*).

The current approaches used to study T cell priming and differentiation *in vivo* depend on TCR transgenic models or identification of epitope-specific T cells by pMHC tetramers (*78*). The *in vitro* differentiation method relies on the use of anti-CD3/CD28 antibodies and exogenous recombinant cytokines (*4, 12, 13*). These three approaches, although very informative, fail to address the diverse specificity of TCRs or the heterogeneity of the responses. Additionally, since specific signals received by a naïve CD4 T cell during a critical window of differentiation are likely to dictate its fate, it is tempting to speculate that cytokine-driven T cell differentiation, although important for the analysis of certain aspects of T cell biology, deviates considerably from a physiological immune response. The *in vitro* priming approach described here enables the study of the antigen-specific CD4 T cell differentiation using physiological levels of stimulation provided by DCs to generate multi-lineage and oligoclonal pathogen-specific T cells. Furthermore, genetic manipulations can be used to explore the roles of molecular pathways in both CD4 T cells and DCs that will open avenues for detailed mechanistic studies.

Approaches using DCs and other APCs to prime antigen-specific CD4 and CD8 T cells have been explored in both human and mouse studies (*15, 79*). However, our studies using the murine system have allowed us to establish the specificity and also reveal the *in vivo* relevance of these *in vitro* primed T cells. Notably, we have used splenic DCs as APCs, which are predominately composed of CD11c^+^CD11b^−^ and CD11c^+^CD11b^+^ populations (*80*). This is currently a limitation of our study and future efforts could focus on using specific DC subpopulations to dissect their impact on dictating transcriptional profile of responding CD4 T cells. Additionally, it would be worthwhile to also use tissue resident or migratory DC populations when investigating CD4 T cell responses to tissue invading or compartmentalized pathogens. Tissue-resident DCs, for example, lamina propria CD103^+^DCs, may generate distinct T cell responses and transcriptional profiles that are relevant to mucosal immunity (*31, 81*).

Taken together, our work highlights a novel workflow for studying pathogen-specific CD4 T cell differentiation. Conceptually, the dataset from this study provides experimental evidence for the importance of dendritic cells in dictating global transcriptional programming of antigen-specific CD4 T cells. Integration of a systems biology approach into this *in vitro* priming system empowers high-throughput analysis of anti-microbial T cell responses to discover novel players (such as caspase-1) in CD4 T cell activation and differentiation.

## Acknowledgements

We thank all members of the Pasare lab for helpful discussions and critical reading of the manuscript, Linley Riediger for mouse colony management and genotyping. We thank the members of Genomics and Microarray Core at UT Southwestern Medical Center for their help with the RNA sequencing experiments. Bret Evers, M.D., Ph.D. kindly performed histopathology analysis and blind scoring of the sections. Russell. E. Vance, Ph.D. at University of California, Berkeley, CA, generously provided Casp1Δ10 mice. Fayyaz S. Sutterwala, M.D., Ph.D., at Cedars Sinai Medical Center kindly provided *Il1b*^−/−^ mice. Vishva Dixit, M.D., kindly provided *Asc*^−/−^ mice.

## Author Contributions

Conceptualization, Y.G., E.K.W., and C.P.; Methodology, Y.G., K.D., A.J., and C.P.; Investigation, Y.G., K.D., and A.J., R.A.I.C.; Formal Analysis, Y.G., I.D. and C.P.; Writing-Original Draft, Y.G. and C.P.; Writing-Review and Editing, Y.G., E.K.W., and C.P.; Resources, I.D. and I.R.; Data Curation, I.D.; Funding Acquisition, C.P.

## Experimental Procedures

### Quantification and Statistical analysis

Statistical analysis performed is indicated in the figure legends, analyzed by GraphPad Prism 7. P values<0.05 were considered statistically significant. Significantly differentially expressed genes in RNA-seq experiments were determined using encoded Matlab or Gene Pattern DESeq2 function described in RNA-seq analysis section. Sample sizes were not predetermined by statistical methods.

### Mice

C57BL/6J (RRID:IMSR_JAX:000664), B6.SJL-*Ptprc*^*a*^ *Pepc*^*b*^/BoyJ (B6.CD45.1; RRID:IMSR_JAX:002014) mice were obtained from Jackson Laboratory and maintained in UT Southwestern mouse breeding core facility. STOCK II17a^tm1.1(icre)Stck^/J (*Il17a-cre*; RRID:IMSR_JAX:016879) and B6.129S4-Ifng^tm3.1Lky^/J ‘GREAT’ (*Ifng-ires-yfp*; RRID:IMSR_JAX:017581) mice were obtained from Jackson Laboratory and bred in-house to B6.Cg-Gt(ROSA)26Sor^tm14(CAG-tdTomato)Hze/^J (*Rosa26-flox-stop-flox-tdTomato*; RRID:IMSR_JAX:007914) mice (a gift from Morrison Laboratory, UT Southwestern). Casp1D10 mice were a kind gift from Drs. Russell Vance and Isabella Rauch ((*55*), University of California, Berkeley). *Asc*^−/−^ mice were provided by Dr. Vishva Dixit (Genentech). *II1b*^−/−^ mice were provided by Dr. Fayyaz S. Sutterwala (Cedars Sinai Medical Center). Unless specified, mice were bred and maintained at the specific pathogen-free facility of UT Southwestern Medical Center, provided with sterilized food and water *ad libitum*. Mice used for infection experiments were kept at a conventional animal facility and provided with non-autoclaved food and water *ad libitum*. Age- and sex-matched mice between 6 and 12 weeks of age were used for all experiments. Both female and male mice were used in experiments. All mouse experiments were performed as per protocols approved by Institutional Animal Care and Use Committee (IACUC) at UT Southwestern Medical Center.

### Bacterial strains

*Listeria monocytogenes* (LM 10403 serotype 1, a gift from Dr. James Forman), *Citrobacter rodentium* (strain ICC168, Nalidixic acid-resistant) and *Staphylococcus aureus* (ATCC-25923) were cultured in agar plate of Brain-Heart Infusion, Luria-Bertani with 30µg/ml nalidixic acid, and Tryptic Soy Broth respectively. A single colony was chosen and secondarily expanded in the respective liquid broth with appropriate antibiotics. For *Listeria monocytogenes* infection, bacteria were grown to log phase (OD600=0.6-1) on the day of infection, extensively washed and resuspended in PBS. Mice were injected intraperitoneally with 2×10^4^ colony forming units (CFU) of *L.monocytogenes*. Tissues were harvested 5 days post infection (p.i.). For *C. rodentium* infection, mice were intragastrically administered 1% sodium bicarbonate and 20-30min later, infected with 5×10^8^ CFU of *C.rodentium*. mLNs were harvested for cell sorting 10 days p.i.

### Isolation of mouse lymphocyte populations

Spleen and lymph nodes were harvested from 6-12 weeks old mice. Single-cell suspension was obtained by dissociation using sterile frosted slides and passing through 70μm cell strainer. Red blood cell lysis was performed as needed. Naïve CD4 T cells were isolated according to MojoSort kit protocol (Biolegend). The purity of naïve CD4 T cells was constantly monitored and maintained at >95%CD4^+^MHC-II^−^CD62L^+^CD44^−^. Splenic dendritic cells were isolated from the spleen of B16-FLT3L melanoma injected mouse. Splenocytes were blocked with Fc block (anti-CD32/CD16) and stained with CD11c-biotin (Biolegend), subsequently with anti-Biotin beads (Miltenyi) and isolated using AutoMacs magnetic selection (Miltenyi). The purity of isolated splenic DCs was maintained at >98% CD11c^+^. For all experiments, dendritic cell donor mice and naïve CD4 T cell donor mice were age- and sex-matched. B cells were isolated from sex- and age-matched naïve mouse spleen by isolating CD19^+^ population using CD19-biotin (BD) and AutoMacs (Miltenyi). Lamina propria lymphocytes (LPL) were isolated as previously described (*31*). All genotypes were co-housed for at least 2 weeks before isolating LPL.

### Pathogen-specific CD4 T cell priming

X-VIVO15 serum-free media (Lonza) was used to avoid T cell activity to bovine serum proteins. In some experiments, 10% complete RPMI media (RPMI1640 media (Hyclone), 10% Fetal Bovine Serum (FCS) (Sigma), L-glutamine, Penicillin-Streptomycin, Sodium Pyruvate, β-mercaptoethanol (Sigma)) was used.

CD11c^+^ dendritic cells were pulsed with pathogen lysate (10μg/ml, dose titrated to induce the maximum response and minimum cell death across the panel) at 1×10^6^/ml for 5 hours and then extensively washed. Naïve T cells were labeled with CFSE (5μM, BioLegend). Dendritic cell and T cells were co-cultured in 1:5 ratio for 5-12 days, depending on the experiments.

For live bacteria stimulation experiment, live bacteria were grown to log phase and extensively washed with cell culture media and used to infect dendritic cells at multiplicity of infection (MOI)=6 for 5 hours and incubated in media with gentamicin (200μg/ml, Life Technologies) for 1 hour. Then DCs were washed extensively and cocultured with naïve CD4 T cells. MOI was determined to match bacteria number with the dose of 10μg/ml lysate.

### Pathogen-specific T cell response recall by B cell-mediated re-stimulation

Pathogen-specific T cells generated using the approach described previously were rested with the provision of a low dose of IL-2 (10 unit/ml, Biolegend) for additional two days until active cytokine production waned. B cells were either pulsed with pathogen lysate or blasted with CpG (The Keck oligonucleotide synthesis facility, Yale University) for 18-24 hours in X-VIVO15 serum-free media, extensively washed and irradiated at the dose of 12 Gy using X-ray irradiator (X-RAD320, Precision X-Ray), and cocultured with T cell in 2:1 ratio. T cell responses were assessed 48 hours later.

### T cell polarization (Cytokine-differentiated T cells)

Tissue culture-treated plates were coated with 5µg/ml of anti-mouse CD3 (Biolegend) and anti-mouse CD28 (Tonbo) for 2-4 hours. 1×10^6^/ml Naïve T cells were polarized for 5 days under Th17 polarization (cdTh17) conditions with 10μg/ml anti-IFNγ (Biolegend), 10μg/ml anti-IL-4 (Biolegend), 20ng/ml IL-6 (Peprotech), 5ng/ml TGFβ1 (Peprotech), 10ng/ml IL-1β (Peprotech) and 20ng/ml IL-23 (Biolegend), or Th1 polarization conditions with 10μg/ml anti-IL-4, 50 unit/ml IL-2 and 10ng/ml IL-12 (Peprotech). For some experiments, polarized cells were removed from all polarizing cytokines and plate-bound anti-CD3/CD28, washed and cultured with 10U/ml IL-2 for an additional 5 days.

### Retroviral Transduction of Th17 cells

Retrovirus was prepared from 1×10^6^ Platinum-E cells transfected with 2.5ug vector and 0.63ug pcl-ECO using Lipofectamine-2000 transfection reagent (Thermo). Viral supernatant was harvested from Platinum-E cultures after 48hrs and 72hrs of transfection. 50U/ml of IL-2 (Biolegend) and 10ug/ml of protamine sulfate (Sigma) was added to virus sup prior to transduction. Naïve CD4 T cells were prepared and differentiated as described before. 24hrs and 48hrs after activation, 1×10^6^ T cells were transduced with 1ml of Virus sup under spin-infection of 2,500 rpm for 90min at 32°C. T cells were returned to the original activation media after spin-infection. 5 days after T cell activation, cells were harvested and performed staining for hCD2 as transduction efficiency marker, as well as intracellular staining.

### T cell adoptive transfer

Pathogen-primed CD4 T cells were extensively washed with plain RPMI1640 media without serum or additives and injected intravenously into the recipient mice (2×10^6^ CD25^+^ICOS^+^ cells/mouse). Recipient mice were co-housed with their respective controls and experimental groups for at least two weeks before the experiment and during the experimental period.

### T cell transfer model of colitis

5×10^5^ CD4^+^CD62L^hi^CD44^lo^CD45RB^hi^CD25^−^ T cells were FACS sorted from spleens and lymph nodes of mice of each genotype, washed and injected intraperitoneally (i.p.) into *Rag1*^−/−^ mice. Weights were measured weekly from 1-3 weeks and biweekly from 4-7 weeks. Serum was collected every week via submandibular bleeding. Mice were sacrificed between 7 and 8 weeks post transfer or when weight loss exceeded 20% of initial weight and colon length was measured at the time of sacrifice. Mice were sacrificed 4-5 weeks post transfer. Colons were removed from the cecum to anus, photographed, fixed with formalin and submitted to the University of Texas Southwestern Molecular Pathology Core for paraffin embedding, sectioning, and H&E staining. Digital images were obtained using Zeiss Axiovert 100 Inverted Microscope with Jenoptik Gryphax Camera. Histology scores were blind-scored by a pathologist (Dr. Bret Evers) at UT Southwestern using the criteria listed in Table S8. LPL cells were isolated from colons of diseased mice following previously established protocols (*67*) and intracellular staining was performed as described in Flow cytometry and FACS section (Supplemental Experimental Procedures).

### Enzyme-linked immunosorbent assay (ELISA)

Briefly, coating antibodies were diluted and coated in ELISA plate overnight at 4°C. Blocked with PBS containing 10%FCS or 1%Bovine Serum Albumin (Sigma). Samples were loaded in duplicates, diluted in blocking buffer and incubated overnight. Detection antibodies were used according to manufacturer’s instruction. Protein concentrations were quantified using TMB or OPD colorimetric assay. Plates were washed extensively in between steps with PBST (Phosphate buffered saline, 0.05% Tween-20).

### Western blot analysis

Cells were extensively washed with PBS on ice and directly lysed in boiled SDS-containing 2X Laemmli buffer. Protein concentration was measured by detergent-resistant Bradford assay. 5-10μg of each lysate was loaded to SDS-Page and immunoblot was performed using standard protocols. Antibodies used for western blot are listed in the resource table.

### Plasmids

MSCV-IRES-hCD2tm* vector was described previously (*82*). Full-length caspase-1 or caspase-1 deficient of CARD was cloned into backbone using XhoI and NotI locus using primers listed in Table S1. Enzymatically inactive caspase-1 was cloned by mutating cysteine (TGC) 284 to alanine (GCT) of the full-length caspase-1.

### RNA isolation and quantification, qRT-PCR

Cells were either sorted into Trizol LS reagent or sorted into complete media and lysed with Trizol reagent. RNA was extracted using the miRNeasy (Qiagen) and treated with DNase I (Qiagen), according to the manufacturer’s instructions. Quantities of extracted RNA were determined using NanoDrop2000 for cDNA synthesis or seq Library preparation. All RNA sent for library preparation qualified for RNA integrity number (RIN)<8.

cDNA synthesis was carried out using M-MLV reverse transcriptase (Invitrogen) in the presence of RNase inhibitor (Promega). Quantitative RT-PCR was performed using SYBR green mastermix (Applied Biosystems) and QuantStudio 7 Flex Real-Time PCR System.

### Data and Software Availability

Illustrations were created with Biorender (https://biorender.io/). Relevant data software packages are listed in Table S2. MatLab code used for RNA-seq analysis is described in ‘RNA-sequencing analysis’ section (Supplemental Experimental Procedures) and is available upon request.

